# Spinal Mechanisms in Post-Activation Potentiation: Facilitation of Presynaptic Inhibition Contrasts H-Reflex Amplitude Reduction

**DOI:** 10.64898/2026.01.16.699288

**Authors:** Miloš Kalc, Aleš Holobar, Matej Kramberger, Nina Murks, Jakob Škarabot

**Author notes:** The authors are co-corresponding authors (Miloš Kalc, Jakob Škarabot).

## Abstract

This study investigated the spinal neural mechanisms underlying post-activation potentiation in ten healthy young males (21.9 ± 4.8 years). Participants performed a 10-second maximal isometric plantarflexion, after which we measured twitch torque and assessed spinal excitability using the soleus H-reflex, D1 presynaptic inhibition and heteronymous Ia facilitation (HF). High-density surface EMG was decomposed to track single motor unit responses. The conditioning contraction increased twitch torque by 12.2 Nm (p < 0.001) immediately and returning to baseline within nine minutes. This mechanical potentiation was accompanied by a 29% reduction in H-reflex amplitude (p < 0.001), which recovered within three minutes. Paradoxically, neurophysiological indices of presynaptic inhibition, D1 and HF were significantly increased (D1: p<0.017; HF: p<0.001), resulting in spinal facilitation. Single MU analysis revealed increased discharge probability, particularly in higher-threshold units indicating overall spinal facilitation. These results demonstrate that post-activation potentiation involves a complex dissociation: H-reflex pathway inhibition along with facilitation of presynaptic spinal mechanisms. This paradox can be explained by either post-activation depression (caused by depletion of neurotransmitter at the Ia–motoneuron synapse) or muscle thixotropy, a contraction history-dependent decrease in muscle spindle sensitivity, which reduces the efficacy of the Ia afferent volley independently of spinal inhibitory mechanisms. Our findings highlight a dissociation between spinal presynaptic facilitation and the decreased H-reflex, underscoring the need for future studies to explicitly test the roles of post-activation depression and muscle thixotropy during post-activation potentiation.

**New & Noteworthy:** This study provides evidence that post-activation potentiation reduces the soleus H-reflex amplitude while concurrently facilitating presynaptic spinal mechanisms. By combining global EMG and single motor unit analyses extracted from high-density surface EMG, we reveal a dissociation between spinal disinhibition and reflex depression. These findings suggest that acute post-contraction reflex suppression might be mediated by mechanisms other than presynaptic inhibition, potentially involving post-activation depression spinal mechanisms or changes in muscle spindle sensitivity.

## Introduction

Muscle contraction history significantly influences subsequent muscle contractions, which can either be enhanced or depressed. For example, electrically elicited tetani or short-lasting (<10 s) voluntary conditioning contractions (Vandervoort et al. 1983) temporarily increase single-twitch performance. This phenomenon, known as post-activation potentiation (PAP) (Sale 2002), is primarily influenced by the intensity and duration of the conditioning contraction (Vandervoort et al. 1983). The principal mechanism behind the observed twitch potentiation is the phosphorylation of the regulatory myosin light chain (Zero and Rice 2021). This phosphorylation alters myosin head orientation, enhancing the likelihood of force-generating cross-bridge interactions by reducing the distance to actin-binding sites (Levine et al. 1996). As a result, Ca²⁺ sensitivity is increased, making twitch force production more efficient, particularly at lower contraction intensities. Indeed, some studies have reported twitch force increases up to 200% immediately following a conditioning contraction (Baudry and Duchateau 2004), with potentiation tapering off exponentially over 10 minutes.

Although the primary mechanisms of PAP are well-established, emerging evidence suggests that neural processes also contribute (Tillin and Bishop 2009; Xenofondos et al. 2015). However, the precise nature of neural contribution is equivocal, with studies showing Hoffmann reflex (H-reflex) depression (lateral gastrocnemius, LG: Güillich and Schmidtbleicher 2006; soleus, SOL: Enoka et al., 1980), enhancement (LG: Güillich and Schmidtbleicher 2006; SOL: Güillich and Schmidtbleicher 2006; Trimble and Harp 1998; vastus medialis: Folland et al. 2008), or no change (Hodgson et al. 2008; Iglesias-Soler et al. 2011; Wallace et al. 2019) following a conditioning contraction (see Zero & Rice, 2021 for a comprehensive review). These discrepancies likely stem from variations in participant characteristics, the nature of the conditioning contractions, time of assessment relative to conditioning contraction, H-reflex assessment methods, muscle selection, and training status (e.g., enhancement more likely shown in trained athletes: Güillich (1996), Folland (2008)).

Because reflex potentiation tends to emerge several minutes after a conditioning contraction, while twitch potentiation typically dissipates more rapidly, some have argued that the delayed reflex effects are not directly related to PAP that mainly contributes to immediate increases in twitch force (Blazevich and Babault 2019). Instead, they may reflect post-activation performance enhancement (PAPE) (Zero and Rice 2021), which typically arises 5 to 10 minutes after a contraction, and encompasses improvements in neuromuscular function not directly linked to myosin regulatory light chain phosphorylation, such as elevated muscle temperature, fluid shifts, or increased blood flow. However, the early H-reflex depression has been linked more closely to PAP and is thought to reflect a compensatory neural strategy, possibly aimed at offsetting the enhanced muscle output (Zero and Rice 2021), but the underlying mechanisms remain experimentally unverified.

A fundamental challenge in interpreting H-reflex changes lies in disentangling presynaptic (e.g., presynaptic inhibition) from postsynaptic (e.g., motoneuron excitability, reciprocal inhibition) mechanisms (Theodosiadou et al. 2022). To address this, electrophysiological techniques often employ conditioning stimuli, such as low-intensity volleys to antagonistic nerves, which elicit D1 (5–30 ms) and D2 (70–200 ms) inhibition responses largely attributed to presynaptic inhibition of Ia terminals (Knikou 2008). However, this approach has known limitations: postsynaptic effects may temporally overlap with presynaptic inhibition, while long-latency cutaneous facilitation or shifts in motor unit recruitment thresholds may confound interpretation by mimicking or masking true presynaptic changes (Hultborn et al. 1987). To resolve these ambiguities, Hultborn and colleagues (1987) demonstrated that, within the first ∼0.6 ms of the H-reflex response, monosynaptic Ia excitation remains uncontaminated by disynaptic inputs, making this early window a reliable indicator of pure Ia excitatory postsynaptic potentials (EPSPs). In principle, the most robust way to distinguish changes in presynaptic inhibition from postsynaptic motoneuron recruitment gain is through peri-stimulus time histogram (PSTH) analysis of single motor unit (MU) discharges, which reflects the direct timing and probability of motoneuron activation in response to synaptic input. Although recent advances in high-density surface electromyography (HDsEMG) decomposition now allow the extraction of single MU responses during evoked contractions (Kalc et al. 2022a, 2022b; Magdič et al. 2025; Škarabot et al. 2023), this approach requires the delivery of a large number of stimuli, typically several tens to construct reliable PSTHs, making it impractical for studies of acute phenomena like PAP, where the time window for capturing spinal adaptations is limited to a few minutes. To address this constraint, we propose to assess the heteronymous Ia femoral facilitation (Grosprêtre et al. 2018; Pierrot-Deseilligny and Mazevet 2000), which in combination with D1 presynaptic inhibition assessment offers a more time-efficient method that allows a robust indirect quantification of presynaptic inhibitory effects on soleus Ia terminals.

In this study, we aimed to investigate the immediate and short-term effects of post-activation potentiation on neuromuscular responses following a maximal voluntary isometric conditioning contraction. Specifically, we examined the twitch force response and H-reflex amplitude, while also exploring potential contributions from presynaptic spinal mechanisms. Furthermore, leveraging a recently developed methodology to extract single MU discharges from HDsEMG signals in evoked contractions (Kalc et al. 2022b, 2022a; Škarabot et al. 2023), we investigated how MU discharge patterns and amplitude cancellation may influence the observed H-reflex responses. Given the involvement of recreationally active participants in this study, we hypothesised that the conditioning contraction would immediately but transiently increase twitch force and concurrently decrease the H-reflex amplitude, with reflex responses returning to baseline within approximately five minutes.

## Methods

### Procedures

#### Participants

Ten young adult males (age 21.9 ± 4.8 years) participated in a single visit counterbalanced crossover repeated measures study. All participants were physically active at least 3-times per week with no history of neurological disorders or major lower-extremity neuromuscular injury as determined by a health history questionnaire (Par-Q; Thomas et al., 1992). Participants were instructed to avoid the consumption of caffeinated beverages and strenuous physical activity 24 hours before the experimental session. Each participant was briefed about any potential risks and discomfort they might face, and they provided written informed consent prior to involvement in the study. This research complied with the most recent principles outlined in the Declaration of Helsinki (except for registration in database), and it received clearance from the Research Ethics Committee of Slovenia (n 0120-84/2020/4).

#### Study design

Participants first visited the laboratory for a familiarisation session, during which they were acquainted with the electrical stimulation paradigms used in the study. They returned three to seven days later for the experimental session. The session began with a standardised warm-up consisting of isometric plantarflexion contractions lasting 4–6 seconds at 50% (2 repetitions), 60% (2x), 70% (1x), 80% (1x), and 90% (1x) of the participant’s maximal perceived effort. This was followed by a force-tracking preliminary procedure (see *Force tracking preliminary procedure*). Subsequently, stimulation parameters for the tibial, common peroneal, and femoral nerves were determined. A 20-minute washout period was then introduced to reduce potential residual effects of the contractions on subsequent measurements.

During the experiment, participants were instructed to remain seated in the ankle dynamometer and perform one of two interventions: (i) a 10-second maximal voluntary isometric contraction (EXP), or (ii) passive rest (CON). Following each intervention, mechanical and electrophysiological responses to stimulation were recorded over an 18-minute period at 1, 3, 5, 7, 9, 12, 15, and 18-minutes post-intervention. This design, omitting baseline assessments prior to the intervention, was adapted from previous work (Folland et al. 2008; Wallace et al. 2019). A 20-minute washout period was included between the two conditions to minimise carryover effects. The warm-up was repeated 10 minutes before the second intervention. Intervention order was counterbalanced across participants to reduce potential order effects. All testing was conducted in a temperature-controlled room maintained between 22 and 24 °C. A schematic representation of the intervention timeline is shown in Figure 1A.

**Figure 1:**
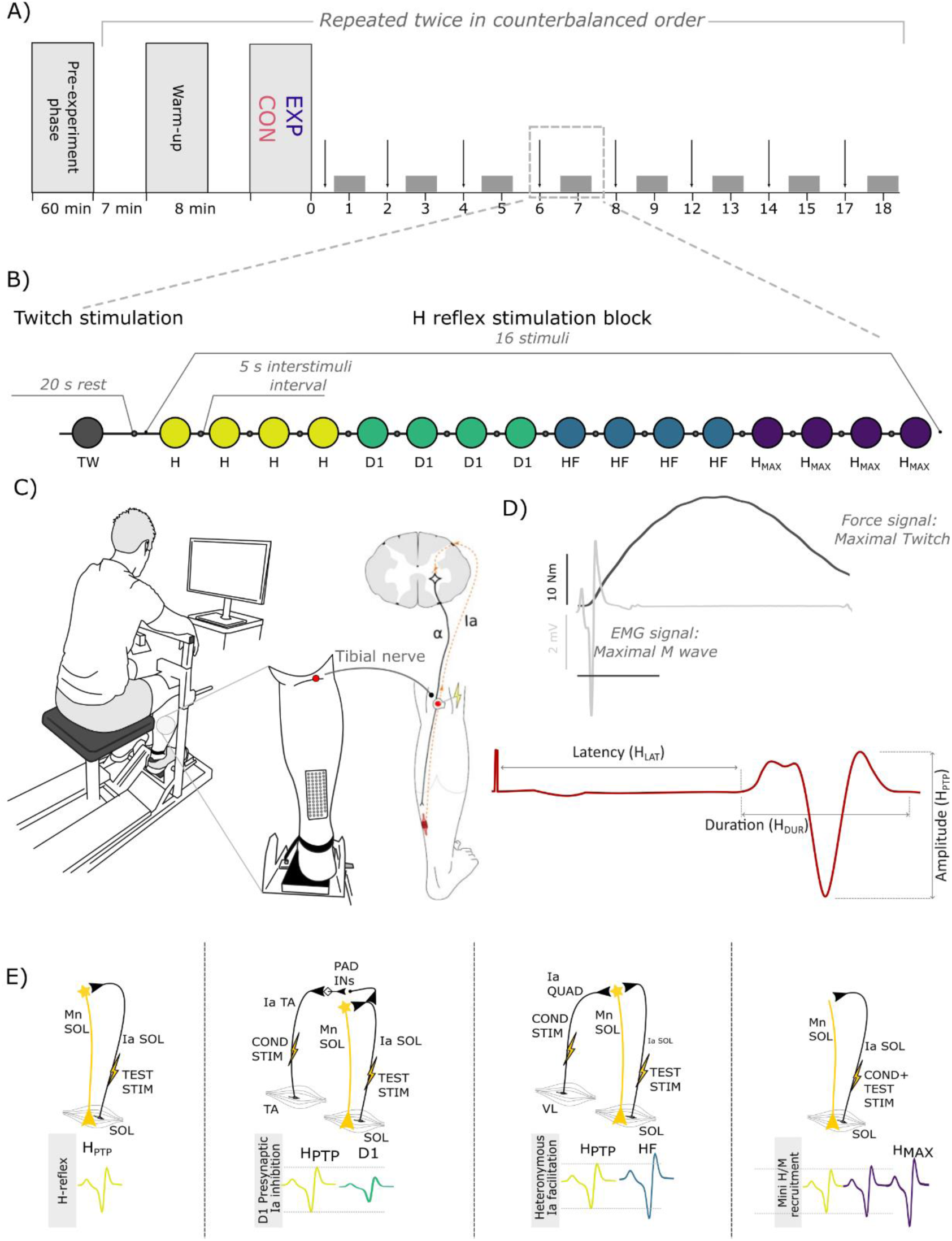
Study design and schematic representation of the assessment procedures. A) Study timeline, including the pre-experiment phase and two experimental conditions (EXP and CON). Vertical arrows indicate the timing of supramaximal electrical stimulations, while the grey boxes represent the blocks of stimulations used to assess spinal reflexes. B) Detailed depiction of the stimulation block, consisting of one supramaximal stimulus (twitch, TW) and 16 submaximal stimuli. These stimuli are grouped in sets of four, designed to assess: H-reflex at the ascending part of the H-M recruitment curve (H), D1 presynaptic Ia inhibition, heteronymous Ia facilitation (HF), and a mini H-M recruitment curve to evaluate the maximal H-reflex. C) Illustration of the participant’s posture on the ankle dynamometer and the points of H-reflex stimulation along the tibial nerve pathway and the placement of the electrode array. D) Representative examples of mechanical (torque) and electrophysiological (M-wave) responses to a single supramaximal stimulus (upper traces), and a typical H-reflex response to submaximal stimulation (lower trace), with key features extracted from the H-reflex waveform (peak-to-peak amplitude, latency, and duration). E) Schematic representations of the spinal circuits involved in different types of reflex modulation (from left to right): H-reflex, D1 presynaptic Ia inhibition, heteronymous Ia facilitation, and maximal H-reflex. The corresponding electrophysiological responses are shown for each reflex type.

#### Participant positioning

Participants were seated upright in the custom ankle dynamometer with their hips, knees, and ankles flexed at 90° (180° = full extension). The right leg was used for all measurements. The foot was strapped securely over the metatarsal region onto the footplate, with the lateral malleolus aligned with the device’s axis of rotation (Figure 1C). A rigid support was placed over the thigh, just above the knee, to limit unwanted movement during plantarflexion. Participants rested their forearms on the device’s arm supports and were instructed to maintain a relaxed posture. To reduce variability in spinal responses, participants were asked to keep their gaze fixed on a monitor in front of them, maintain a neutral head position, and avoid clenching their jaw or squeezing their hands during stimulation (Mitsuyama et al. 2017).

### Force tracking preliminary procedure

Following the warm-up, participants rested for 60 seconds before performing two maximal voluntary isometric contractions (MVCs), separated by 120 seconds of rest. They were instructed to gradually increase force to maximum over 1–2 seconds and maintain maximal effort for approximately 3 seconds. Strong verbal encouragement was provided to ensure maximal effort (Verges et al. 2009). After MVC determination, participants completed submaximal isometric contractions lasting 20 seconds at 10%, 20%, and 30% MVC, and 15 seconds at 40%, 50%, and 70% MVC. Each contraction was followed by a 120-second rest period to prevent fatigue. These force-tracking contractions were used to establish MU separation filters, facilitating MU identification during electrically elicited contractions (Kalc et al. 2022a; Škarabot et al. 2023), see ‘HDsEMG processing’ section for details.

### Electromyographic recordings

Electromyographic signals were recorded from the soleus (SOL) muscle using both high-density surface EMG and bipolar EMG (biEMG). Prior to electrode placement, the skin over the SOL region was shaved and abraded with abrasive paste (Piervirgili et al. 2014). The electrode array was mounted on the muscle belly covering the central portion of the soleus muscle, with the long side of the array aligned with the longitudinal axis of the muscle (Figure 1C). This electrode position has been suggested as the position where the H reflex with the largest amplitude can be observed on the soleus muscle (Botter et al. 2015). The biEMG electrodes were positioned laterally to the array, with a reference electrode positioned over the tibial tuberosity, and a ground electrode wrapped around the ankle. The HDsEMG was used to extract individual motor unit spike trains from both voluntary and evoked contractions, whereas the biEMG setup allowed for real-time monitoring of short signal segments (30 ms before to 70 ms after each electrical stimulus), ensuring visual feedback of the evoked responses and signal quality.

### Posterior tibial nerve stimulation

H-reflexes, M-waves, and mechanical twitches were elicited in the right soleus (SOL) muscle using single rectangular pulses (1 ms duration) delivered to the posterior tibial nerve. The stimulation anode was placed over the patella, while the optimal site for cathodal stimulation in the popliteal fossa was first identified using a stimulation pen and then marked for electrode placement.

To obtain the H-reflex and M-wave (HM) recruitment curve, stimulation began at 5 mA and was progressively increased in 1 mA steps until the maximal H-reflex amplitude (H_MAX_) was reached. Once the H-reflex began to decline, stimulation intensity was further increased in 5 mA increments until no further increase in M-wave amplitude was observed. To ensure supramaximal stimulation, the final intensity was increased by 50%. These supramaximal stimuli were used to evoke maximal twitch peak torque (PT) and maximal M-wave (H_MAX_).

To investigate the different spinal pathways after the interventions, the H-reflex was conditioned with nerve stimulations of homonymous spinal pathways to assess changes in the level of presynaptic inhibition acting on Ia afferent terminals. Unconditioned H-reflexes served as a baseline for conditioned volleys arising from i) stimulation of the common peroneal nerve to elicit presynaptic inhibition (D1) and ii) stimulation of the heteronymous nerve (femoral nerve) to elicit heteronymous Ia facilitation pathways (HF). Additionally, a mini recruitment curve was used to determine changes in maximal H-reflex post-intervention. A block of 16 stimuli (Figure 1B), comprising four unconditioned H-reflexes, four stimuli each for assessing D1 and HF, and four with progressively increasing intensity for H_MAX_, were delivered at 6-second interstimulus intervals (Figure 1E).

### Unconditioned H-reflex

To evoke the unconditioned SOL H-reflexes, the tibial nerve was stimulated with an intensity that induced a response of the SOL H-reflex in the ascending part of the HM recruitment curve preceded by a visible small M-wave (M_atH_) of amplitude between 5 and 10% of M_MAX_ (Knikou 2008). The reproducibility of M_atH_ was used to monitor the stability and reliability of the stimulation. A stable M_atH_ indicates that a constant number of motor nerve fibres and Ia afferents are excited by the same stimuli, ensuring the consistency of stimulus condition within the experimental session (Grosprêtre et al. 2014). Thus, the intensity was slightly adjusted during the experiment to elicit a consistent M_atH_ amplitude. The peak-to-peak amplitude of the unconditioned H-reflex (H_PTP_) was further considered in data analysis.

### Assessment of D1 presynaptic inhibition

H-reflexes were conditioned to induce D1 presynaptic inhibition of Ia afferents onto α-motoneurons, following the methodology presented by Knikou (2008). Conditioning stimuli, consisting of triplets at 300 Hz (1 ms width), were delivered to the branch of the common peroneal nerve activating ipsilateral pretibial flexors. Electrical stimulation was delivered with an intensity set at 1.2 times the motor threshold of the tibialis anterior (TA; Achache et al., 2010). Optimal conditioning-test timing for maximal H-reflex inhibition was determined for each participant at the start of the session, exploring intervals from 25 ms to 10 ms in 1 ms steps. The timing producing the highest inhibition was used. The ratio between the conditioned (D1) and the unconditioned reflex (H_PTP_) indicates the activation of presynaptic inhibitory mechanisms (see Figure 1D) and was further considered in data analysis.

### Assessment of Heteronymous Ia Facilitation

To examine HF, a conditioning stimulus was applied to the ipsilateral femoral nerve (Figure 1E). Single stimulations (1 ms pulse width) were set at 1.3 times the motor threshold of the vastus lateralis muscle. The most effective conditioning test interval for facilitating soleus HF amplitude was identified early in the experimental session, with intervals ranging from -10 ms to +5 ms in 1 ms steps. The timing that produced the highest facilitation in each participant was used. Special attention was given to the conditioning-test interval to ensure monosynaptic interaction of quadriceps Ia afferents with the triceps surae motoneuronal pool (Morita et al. 1995), as facilitations at larger intervals might also involve polysynaptic circuits. Bergmans et al. (1978), have noted that heteronymous Ia facilitation of the H-reflex is not always observable in the lower limbs of all participants. Therefore, only participants who exhibited an increase in SOL H-reflex after a heteronymous conditioning impulse to the femoral nerve in familiarisation were invited to participate in the experimental session. The ratio between the conditioned (HF) and the unconditioned reflex (H_PTP_) indicates the facilitation of the SOL Ia–α motoneuron after femoral nerve activation (Hultborn et al., 1987; Figure 1D) and was further considered in data analysis.

### Maximal H-reflex amplitude

The maximal H-reflex was evoked by four stimuli starting at an intensity of 2 mA below the H_MAX_ reached at the beginning of the visit (1 mA increments). The H-reflex response that produced the highest peak-to-peak amplitude of the series of 4 stimuli was considered H_MAX_ for that time point.

### Stimulation protocol justification

The stimulation protocol for studying spinal mechanisms in this study involved balancing the need for a sufficiently long inter-stimuli interval to prevent homosynaptic depression (Özyurt et al. 2020) in a resting muscle and the need for a significant number of stimuli to counteract the intrinsic variability of H-reflex amplitude (Gossard et al. 1994). Though longer intervals (∼10 s) are crucial to reduce homosynaptic depression risk (Hultborn et al. 1996; Özyurt et al. 2020; Stein et al. 2007), and 5 to 10 stimuli are typically needed to obtain a reliable measure of the physiological response (Theodosiadou et al. 2022), we aimed to record as many responses as feasible within a limited timeframe (90 s), targeting 16 stimuli divided into four groups. During the familiarization session, we experimented with shortening the inter-stimuli interval to determine the shortest possible duration without inducing homosynaptic depression. A 6-second interval emerged as the optimal compromise, as the shortest duration not triggering homosynaptic depression in our participants, thereby allowing for a greater number of stimuli within the given timeframe.

### Instrumentation

#### Isometric dynamometer

Isometric plantarflexion torque was recorded using a custom-made ankle dynamometer (Wise Technologies, Ljubljana, Slovenia). A load cell (Z6FC3/200kg, AEP transducers, Modena, Italy) was embedded in the footplate to measure plantarflexion force.

#### Electromyography systems

HDsEMG signals were recorded using a semi-disposable adhesive electrode array with 5 columns × 13 rows and 8 mm interelectrode distance (GR08MM1305, OT Bioelettronica, Torino, Italy). Signals were amplified with a 16-bit amplifier (Quattrocento, OT Bioelettronica) and acquired at 5,120 Hz using OTBiolab+ software. A reference electrode (5 × 3 cm, T3545) was placed on the tibial tuberosity, and grounding was achieved using a water-soaked strap (WS2, OT Bioelettronica, Torino, Italy) around the ankle. Signals were band-pass filtered between 10–500 Hz.

Bipolar EMG signals were recorded with surface electrodes (Covidien 24 mm, interelectrode distance 25 mm) placed lateral to the HDsEMG array. The reference electrode (50 × 100 mm, 00734, Compex, UK) was placed over the patella. Signals were sampled at 10,000 Hz using PowerLab hardware and LabChart software (ADInstruments, Australia) and band-pass filtered (10–500 Hz).

#### Electrical stimulation equipment

The posterior tibial nerve was stimulated using a custom build constant-current stimulator (Stim_1, EMF Furlan, Ljubljana, Slovenia) delivering 1 ms rectangular pulses. The anode (50 × 90 mm, MyoTrode PLUS, Globus, Italy) was placed over the patella, and the cathode (25 mm diameter, J10R00, Axelgaard) over the popliteal fossa.

The common peroneal and femoral nerves were stimulated using the Digitimer DS7R stimulator (Digitimer, UK). Two silver chloride electrodes (8 mm) were placed over the upper anterolateral leg, distal to the fibular head to stimulate the common peroneal nerve. For femoral nerve stimulation, the cathode (25 mm diameter, J10R00, Axelgaard Manufacturing Co., Lystrup, Denmark) was placed in the femoral triangle and the anode (50 × 90 mm, MyoTrode PLUS, Globus, Italy) under the gluteus maximus. A manual switch was used to alternate stimulation targets.

### Data processing

#### Global EMG

EMG reflex responses were extracted from the array electrode. Two sets of five neighbouring monopolar signals within the central portion of the HDsEMG (columns 2–4 and rows 4–7 of the bidimensional array) were averaged and differentiated to obtain a bipolar EMG derivation with an equivalent interelectrode distance of 1.6 cm (Del Vecchio et al. 2017, 2018). Although EMG measurements were also taken in a bipolar configuration, we chose to extract the data from the array electrode because the H-reflex amplitude can vary substantially depending on the recording electrode’s position (Botter et al. 2015), and because we aimed to obtain both global EMG and decomposed EMG from the same source.

Maximal M peak-to-peak amplitudes (M_MAX_) were extracted from supramaximal stimulations. Following recent H-reflex reporting guidelines by Theodosiadou et al. (2022), various parameters were extracted from the reflex electrophysiological response, including normalized peak-to-peak amplitude (H_PTP_/M_MAX_), reflex latency (H_LAT_), and duration (H_DUR_) (Figure 1D). Additionally, to study D1 presynaptic inhibition and HF, we calculated the amplitude ratios of D1 and HF conditioned reflexes to their respective unconditioned responses: D1/H_PTP_ and HF/H_PTP_.

#### Torque data

Torque signals were filtered with a low-pass fourth-order zero-lag Butterworth filter at a cutoff frequency of 25 Hz. From these filtered signals, the peak twitch torque (TW_PT_) was calculated for the supramaximal stimulations.

### HDsEMG processing

An extended explanation of the procedure involved in the detection of single MUs discharges during elicited contractions can be found in our previous works (Kalc et al. 2022b, 2022a; Škarabot et al. 2023). Briefly, HDsEMG signals were decomposed using the well-established Convolution Kernel Compensation (CKC) algorithm (Holobar and Zazula 2007). First, data from isometric force tracking contractions collected at the beginning of the experimental session (see ‘*Force tracking preliminary procedure*’) were decomposed for each contraction level in isolation (Figure 2A and 2B). The decomposition’s effectiveness was quantitatively evaluated using the Pulse-to-Noise Ratio (PNR), which is a reliable indicator of the motor unit (MU) identification precision (Holobar et al. 2009). Any MUs with a PNR below 28 dB were excluded from further analysis. The firing patterns of the remaining MUs underwent meticulous scrutiny and refinement by an expert. This editing process involved discarding MUs that exhibited either unusually low average firing rates (<8 Hz) or highly variable firing patterns, as indicated by a coefficient of variation of the interspike interval exceeding 0.4 (Drost et al. 2006). For each MU that passed these criteria, its respective MU filter was determined. This filter denotes the specific linear combination of spatio-temporal HDsEMG channel data that approximates the spike train of an individual MU (Farina et al. 2008b; Holobar and Zazula 2007). Notably, in situations involving isometric muscle contractions, an MU filter derived from signals of one contraction can be applied effectively to signals of another contraction of the same muscle to ascertain the corresponding MU spike train (Farina et al. 2008b; Holobar and Zazula 2007). In our experiment, we initially established MU filters from voluntary contraction HDsEMG signals and then applied these filters to the HDsEMG signals of elicited H-reflexes. To avoid redundancy in MU identification across different levels of voluntary contraction, the firing patterns of MUs were compared against each other. Any duplicates identified (i.e., MUs sharing more than 30% of their firings), were removed by retaining only the ones with the highest PNR, thereby ensuring the uniqueness of each MU in the analysis. Subsequently, these refined MU filters were applied to the HDsEMG signals from H-reflexes, facilitating the identification of MU spike trains in evoked contractions (Figure 2C). The identified MU spike trains were segmented into actual MU discharges or baseline noise, an approach inherent to the CKC method (Holobar and Zazula 2007). The final step involved a meticulous manual review by an expert, who compared the presence of crosstalk in MU spike trains derived from both voluntary and evoked contractions, making manual adjustments to the segmentation results where necessary, as suggested previously (Farina et al. 2008b, 2010; Holobar et al. 2009; Negro et al. 2016). It is important to note that only MUs identified in the isometric contractions could be detected in electrically evoked contractions. Moreover, recruitment threshold was estimated as the instantaneous force at the instant of the first spike in the binary spike train.

**Figure 2:**
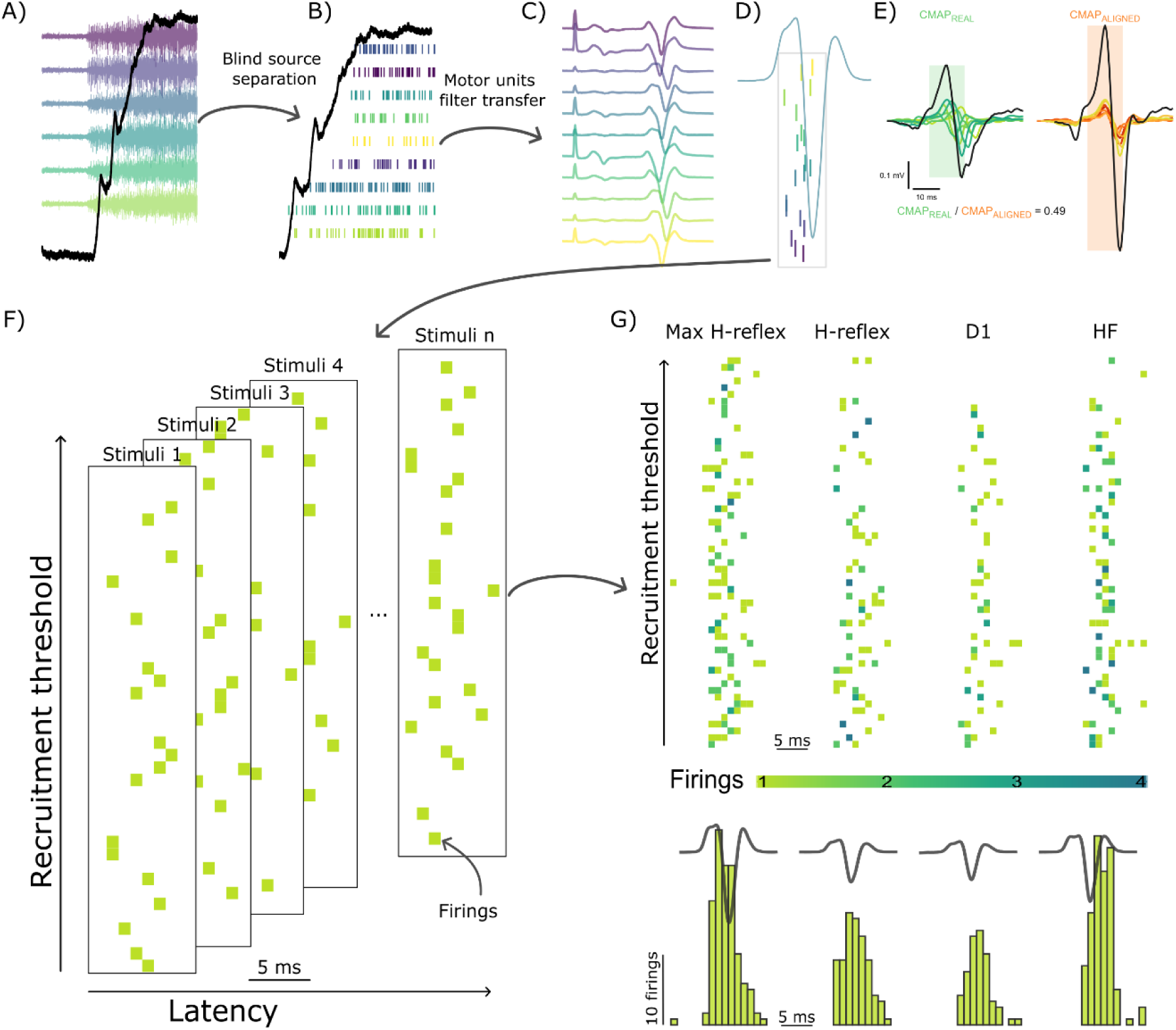
Representative data from a participant and methodological steps to extract single motor unit firings from the HDsEMG recordings. A) Raw electrophysiological signal during a voluntary isometric contraction. B) Extracted motor unit firings during voluntary isometric contractions. C) Reflex response propagation on a sample of channels on an electrode array; D) MU firings extracted form a reflex response. E) Example of CMAP amplitude cancellation. Arithmetic summation of eight MUAPs using recorded latencies and aligned based on the positive peak amplitude, respectively. F) Graphical representation of MU firings extracted from the electrophysiological response in consecutive stimuli. Each panel represent the temporal distribution (latency) of MUs firings with different recruitment thresholds. G) Graphical representation of summated MU firings for the maximal H-reflex, H-reflex, D1 and HR responses, respectively. In the upper figure, the coloured squares represent firings latencies of motor units of different recruitment thresholds. Darker colour represents a higher firing count at same latency at a millisecond precision. The sum of same data is depicted in the histogram at the bottom with the respective global EMG reflex response. Note that the relationship between latencies and recruitment thresholds on the level of individual motor units might not be related, likely due to detection delays (Škarabot et al. 2023); however, at the level of the whole sample higher threshold MUs exhibited longer discharge latencies compared to lower-threshold MUs (see Results).

Following MUs identification, we extracted the MU firing Latencies (HMU_LAT_, D1MU_LAT_, HFMU_LAT_, H_MAX_MU_LAT_) and probability of MU discharge (HMU_PROB,_ D1MU_PROB,_ HFMU_PROB,_ H_MAX_MU_PROB_).

MU discharge ratio was calculated between recognized and discharged MUs for conditioning responses. D1_TO_HMU_RATIO_ and HF_TO_HMU_RATIO_ were calculated subtracting the unconditioned H reflex discharge ratio from D1 and HF discharge ratios, respectively. This metric was calculated to compare single MU data to D1/H_PTP_ and HF/H_PTP_ extracted from global EMG.

In addition, electrically elicited responses generate MU discharges that are highly synchronous; however, axonal conduction velocities, recruitment thresholds, and the spatial distribution of innervation zones introduce small but systematic differences in MUAP latency (Škarabot et al. 2023). MU synchronisation was thus quantified using two complementary measures: (1) the standard deviation of MU firing latencies, and (2) compound motor unit action potential (CMAP) amplitude cancellation. The temporal dispersion of MU discharges was quantified as the *standard deviation* of MU firing latencies for each reflex response. This measure was computed for all evoked responses (HMU_SD_, D1MU_SD_, HFMU_SD_, H_max_MU_SD_) and reflects the spread of MU discharges around the mean response latency. CMAP amplitude cancellation arises from the algebraic summation of the positive and negative phases of MUAPs (Farina et al. 2008a) for each reflex response. When MU discharges occur with temporal dispersion, simultaneous summation becomes less constructive, and the resulting compound waveform exhibits a reduced amplitude. Thus, greater cancellation indicates lower synchrony, whereas lower cancellation reflects highly synchronous MU firing (Figure 2E). To quantify cancellation, we computed two versions of the CMAP for each reflex response: CMAP_REAL_ - the summation of MUAPs obtained using the actual MU discharge latencies, and CMAP_ALIGNED_ - the summation of MUAPs obtained after temporal alignment of all MUAPs at their positive peak, creating an idealised condition of zero temporal dispersion and therefore minimal cancellation. The amplitude cancellation index was then defined in Equation 1:

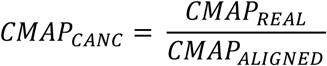

Equation 1: CMAPCANC represents CMAP amplitude cancellation, calculated as the ration between CMAPREAL - the arithmetic summation of MUAPS aligned using their true discharging latencies and CMAPALIGNED the summation of the same MUPAS aligned at their peak values.

Importantly, although MUAP shape strongly influences amplitude cancellation under typical circumstances (Farina et al. 2008a), this does not pose a methodological limitation here because both MUAP_REAL_ and MUAP_ALIGNED_ are constructed from the same set of MUAPs; therefore, any shape-related factor cancels out, and the ratio isolates only temporal dispersion. However, other limitations of CMAP amplitude cancellation-based metrics may bias the results. Indeed, CMAP amplitude cancellation depends on the number of MU discharges (Farina et al. 2008a). This phenomenon particularly affects the results of this study, because cancellation becomes ill-defined when only one or very few MU discharges are present in a single reflex response, causing the cancellation to approach higher values. We addressed this issue by analysing the impact of the number of discharging MUs on CMAP amplitude cancellation and reporting our mitigation procedure in the Supplementary Material (S1).

Please note that the per-MU analyses served as a complementary approach to our primary global EMG analysis. Because meaningful changes in the global EMG were observed only in the early post-intervention period, MU-level analyses were restricted to the first four time points.

### Statistical analysis

Statistical analyses were performed using the R programming language (v.4.2.1; R Foundation for Statistical Computing, Vienna, Austria) within the RStudio environment (2025.09.2). Linear and generalized linear mixed-effects models were used to evaluate changes in neurophysiological and mechanical outcomes, with *lme4* (Bates et al. 2014) and *lmerTest* (Kuznetsova et al. 2017) packages.

To assess the effect of intervention (EXP vs. CON) and timepoint (eight levels) on torque (TW_PT_), a set of global EMG outcomes (peak-to-peak amplitude, latency, and duration of reflex responses), and a set of outcomes calculated from the combined discharges of all MUs within each evoked response (MU discharge ratio, MU discharge standard deviation and CMAP amplitude cancellation), linear mixed-effects models were fitted with *intervention*, *timepoint*, and their interaction as fixed effects. Participant ID were included as random intercepts, with intervention also modelled as a random slope:

*Outcome ∼ Intervention × Timepoint + (1 + Intervention ∣Participant ID)*

MU-level data were extracted only from the first four timepoints. For MU outcomes based on individual firings (MU discharge probabilities and MU latencies), models were extended to include standardised MU R*ecruitment Threshold* as a continuous covariate. Two-way interactions between R*ecruitment Threshold* and both *intervention* and *timepoint* were included:

*Outcome* ∼ *Intervention* × *Timepoint* + *Recruitment Threshold* : *Intervention* + *Recruitment Threshold* : *Timepoint* + (1 + *Intervention* ∣ *Participant ID*)

MU discharge probability, a binary outcome (1 = MU discharged; 0 = MU not discharged), was analysed using a generalized linear mixed model (GLMM) with a *binomial* distribution and *logit* link function in the *lme4* package. Although a logistic link was used, estimated marginal means are reported as predicted probabilities, not odds ratios, to aid interpretation.

Model assumptions (e.g., normality of residuals and random effects, homogeneity of variance, linearity, and multicollinearity) were verified using the *performance* package (Lüdecke et al. 2021). All models were analysed separately for each elicited response condition: H, H_MAX_, D1 and HF.

Post hoc interaction contrasts and simple effects were tested using the *emmeans* package (Lenth et al. 2020), with p-values adjusted using the *multivariate t* method (Genz and Bretz 2002). For torque and global EMG data, the 8th timepoint was used as the reference level for comparisons; for MU-based data, the 4th timepoint served as the reference. Estimated marginal means and 95% confidence intervals were reported for all outcomes, while trends of the continuous covariate are presented graphically. The significance threshold for all analyses was set at α = 0.05. Cohen’s *d* effect sizes were calculated as the difference between estimated marginal means divided by the residual standard deviation of the fitted model. Effect size thresholds were interpreted as: *d* < 0.2 negligible, *d* < 0.5 small, *d* < 0.8 medium, and *d* ≥ 0.8 large (Cohen 1988).

## Results

### Torque results

TW_PT_ was significantly affected by *time × intervention* interaction (F_7, 146.07_ = 27.21; p < 0.001), with interaction contrasts revealing differential modulation of TW_PT_ between interventions at 1, 3, 5 and 7 compared to 18 minutes after contraction (p < 0.001; d = 4.89; p < 0.001, d = 2.01; p < 0.001, d = 1.85; p = 0.005, d = 1.41, respectively **Error! Reference source not found.**C). Simple contrasts revealed that TW_PT_ in the EXP condition were 12.2 Nm (p < 0.001, d = 4.71 ), 4.73 Nm (p = 0.002, d = 1.83), 4.30 Nm (p = 0.004, d = 1.66) and 3.18 Nm (p = 0.028, d = 1.23) higher compared to CON conditions at 1, 3, 5, and 7 minutes after intervention, respectively.

M_MAX_ was not significantly affected by the interaction between *time × intervention* (F_7, 135_ = 1.50, p = 0.169, Figure 3D).

**Figure 3:**
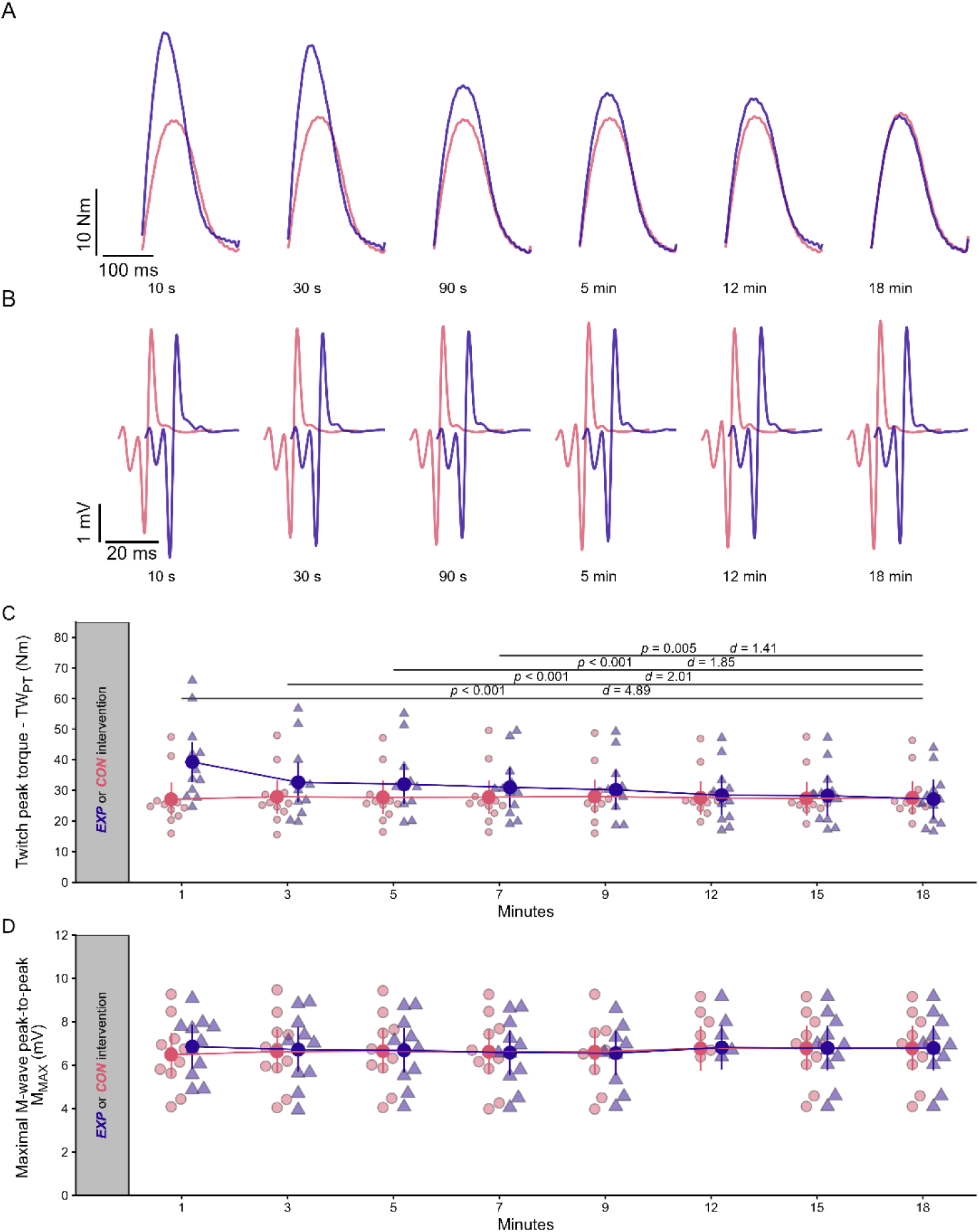
Mechanical and electrophysiological responses to supramaximal electrical stimulations. A) Torque responses over time from a representative participant, illustrating the modulation of torque traces across different time points. B) Representative M-wave responses over time. To facilitate comparison between conditions, traces from the EXP condition are time-shifted by 15 ms to avoid overlap. C) Maximal twitch peak torque across time points. D) Peak-to-peak M-wave amplitudes (MMAX). Statistical data are presented as estimated marginal means with 95% confidence intervals for both the control (CON, red circles) and experimental (EXP, blue triangles) conditions. Smaller and lighter colour circles and triangles represent individual raw data points. Horizontal lines represent statistically significant interaction effects, accompanied by p values and Cohen d values above the relevant lines.

## Global EMG results

H_PTP_/M_MAX_ was significantly affected by the interaction between *time × intervention* (F_7, 496.03_ = 3.38, p < 0.003), with interaction contrasts revealing modulation of H_PTP_/M_MAX_ between interventions at 1 compared to 18 minutes after contraction (p = 0.002; d = 3.60; Figure 4 Figure 4D). Simple contrasts revealed that in H_PTP_/M_MAX_ in the EXP condition was 29.3 percentage points (p < 0.001, d = 3.13) lower compared to CON conditions at 1 minutes after contraction.

**Figure 4:**
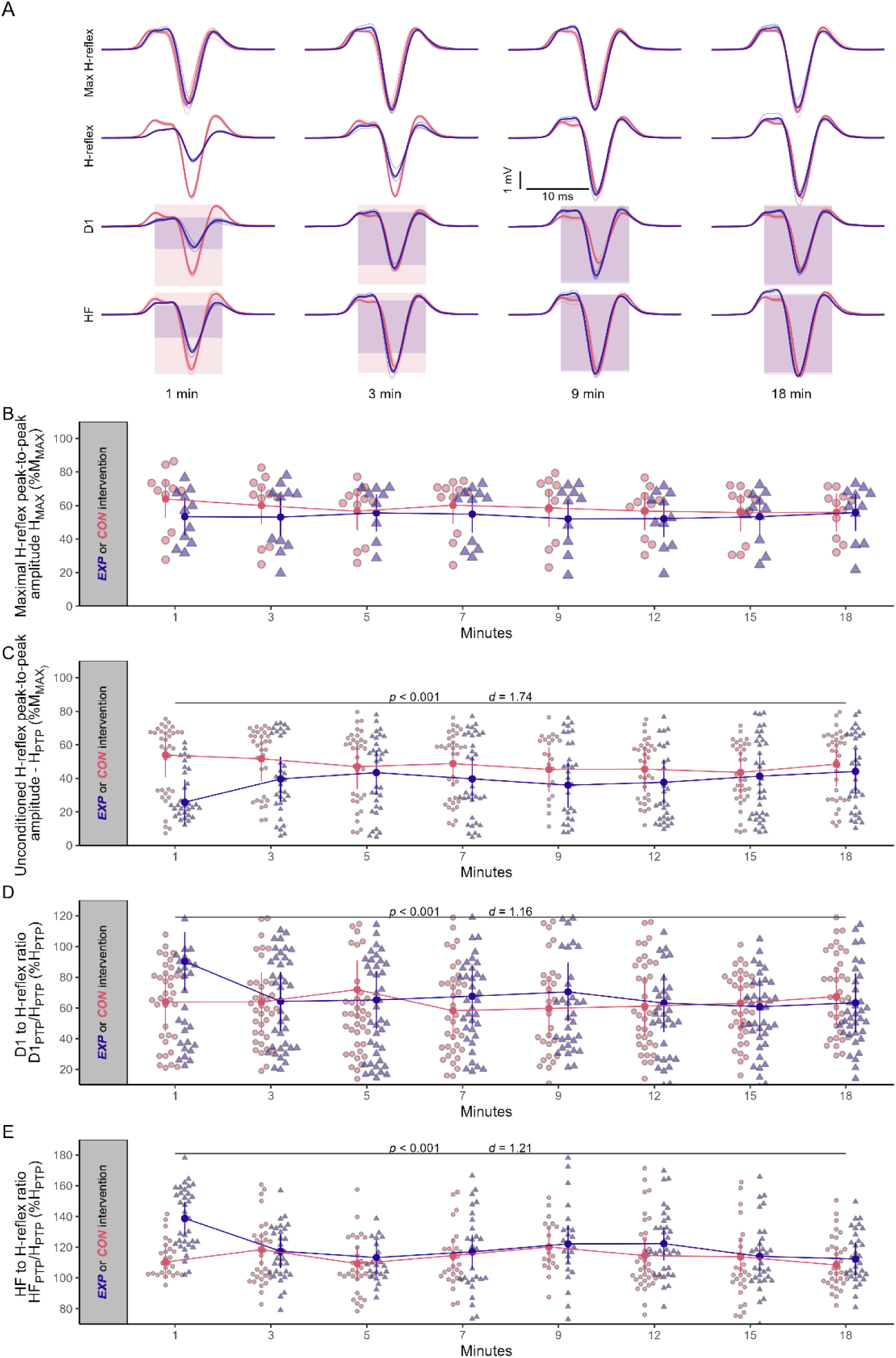
Global electrophysiological responses to submaximal electrical stimulations. A) Unconditioned (HPTP, HMAX) and conditioned (HD1, HHF) H-reflex responses over time from a representative participant, illustrating the modulation of electrophysiological responses across different time points. Thin line traces represent raw signals; meanwhile thick traces represent the mean response. Blue and red box in D1 and HF traces represent the amplitude of the unconditioned H-reflex in the same stimulation block for EXP and CON interventions, respectively, allowing for a visual comparison of the unconditioned and conditioned H-reflex amplitude. B) Maximal H-reflex peak-to-peak amplitude (HMAX) across time points. C) Unconditioned peak-to-peak amplitude of the H-reflex in the ascending part of the H/M recruitment curve (HPTP). D) Ratio between the conditioned (D1PTP) and unconditioned H-reflex (HPTP) peak-to-peak amplitudes. E) Ratio between the conditioned (HFPTP) and unconditioned H-reflex (HPTP) peak-to-peak amplitudes. Statistical data are presented as estimated marginal means with 95% confidence intervals for both the control (CON, red circles) and experimental (EXP, blue triangles) conditions. Smaller and lighter colour circles and triangles represent individual raw data points in the background. Horizontal lines represent statistically significant interaction effects, accompanied by p values and Cohen d values above the relevant lines.

D1/H_PTP_ and HF/H_PTP_ were significantly affected by the interaction between *time × intervention* (F_7, 567.13_ = 3.70, p < 0.001 and F_7, 437.33_ = 4.99, p < 0.001, respectively), with interaction contrasts revealing modulation in both variables between interventions at 1 compared to 18 minutes after contraction (p = 0.017, d = 0.71, Figure 4D; and p < 0.001, d = 1.15, Figure 4E, respectively). Simple contrasts revealed that in D1/H_PTP_ and HF/H_PTP_ in the EXP condition were 17.1 (p < 0.017, d = 0.57) and 28 percentage points (p < 0.001, d = 1.25) higher compared to CON conditions 1 minute after intervention. CON condition remained statistically unchanged for the whole duration of the experiment (p ≥ 0.154).

H_LAT_, H_DUR_, H_MAX_, were not significantly affected by the interaction between *time × intervention* (H_LAT_: F_6,18.12_ = 0.95, p = 0.481; H_DUR_: F_7, 64.82_ = 0.38, p = 0.923; H_MAX_: F_7, 136.00_ = 1.45, p = 0.190, Figure 4B; respectively).

### Single motor unit behaviour results

A total of 537 MUs resulting in 53.7 ± 16.9 (range 21-78) unique MUs per participant were detected (Table 1). The latencies of MU discharges were not significantly affected by the interaction between *time × intervention* in any of the assessed conditioning protocols (HMU_LAT_: F_45112.7_ = 0.99, p = 0.396; D1MU_LAT_: F_3, 4573.3_ = 1.31, p = 0.267; HFMU_LAT_: F_3, 5660_ = 0.48, p = 0.692; H_MAX_MU_LAT_: F_3, 6428.6_ = 1.02, p = 0.378; Figure 5B). However, MU discharge latencies were strongly dependent on MU *recruitment threshold* (HMU_LAT_: F_1, 5117_ = 21.1, p < 0.001; D1MU_LAT_: F_1, 4547.3_ = 10.49, p = 0.001; HFMU_LAT_: F_1, 5655_ = 34.06, p < 0.001; H_MAX_MU_LAT_: F_1, 6426.5_ = 29.22, p < 0.001), with higher-threshold MUs exhibiting longer latencies (Figure 5C). Moreover, a significant *intervention × recruitment threshold* interaction emerged for D1MU_LAT_ (F_1, 4392.3_ = 4.23, p = 0.039), where the EXP condition showed a reduced influence of recruitment threshold on MU firing latency.

**Figure 5:**
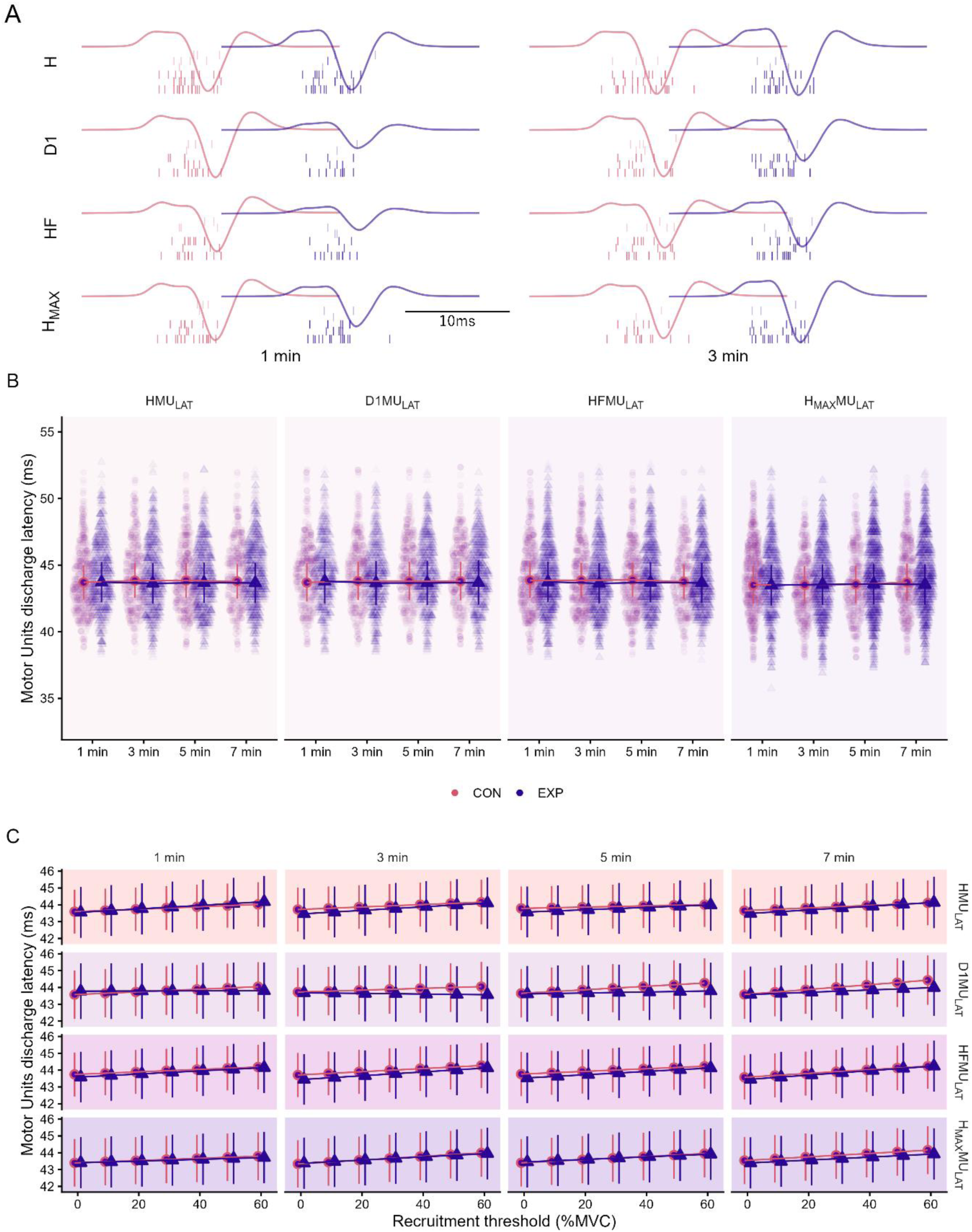
Motor unit (MU) discharges in elicited contractions. A) Unconditioned (H, HMAX) and conditioned (D1, HF) H-reflex responses between 1 and 3 minutes after intervention and respective MU firings from a representative participant. EXP (blue) electrophysiological responses are shifted by 15 ms compared to CON (red) to avoid overlap. Vertical lines represent MU firings. B) MU latency across timepoints. C) Graphical representation of the influence of recruitment threshold on MU latency across timepoint and different stimulation conditions. Statistical data are presented as estimated marginal means with 95% confidence intervals for both the control (CON, red circles) and experimental (EXP, blue triangles) conditions. Smaller and lighter colour circles and triangles represent individual raw data points in the background. Horizontal lines represent statistically significant interaction effects, accompanied by p values and Cohen d values above the relevant lines.

**Table 1:** Descriptive statistics of recognized and discharged Motor Units.

| | n | Mean ± SD / person [range] |
| --- | --- | --- |
| Recognised MUs | 556 | $55.6 \pm 16.6$ [21 - 78] |
| Discharged MUs / response | | $21.8 \pm 13.9$ [0 - 64] |

MU discharge probability showed a significant interaction between *time × intervention* in all assessed conditioning protocols (HMU_PROB_: χ²_3_ = 38.09, p < 0.001; D1MU_PROB_: χ²_3_ = 27.15, p < 0.001; HFMU_PROB_: χ²_3_ = 57.80, p < 0.001; H_max_MU_PROB_: χ²_3_ = 50.04, p < 0.001; Figure 6A). Moreover, a significant *intervention* × *recruitment threshold* interaction emerged for all conditioning protocols (HMU_PROB_: χ²_4_ = 206.80, p < 0.001; D1MU_PROB_: χ²_4_ = 214.267, p < 0.001; HFMU_PROB_: χ²_4_ = 152.67, p < 0.001; H_max_MU_PROB_: χ²_4_ = 115.13, p < 0.001), showing higher probability of MU discharge in higher-threshold MUs in the EXP condition (Figure 6B).

**Figure 6:**
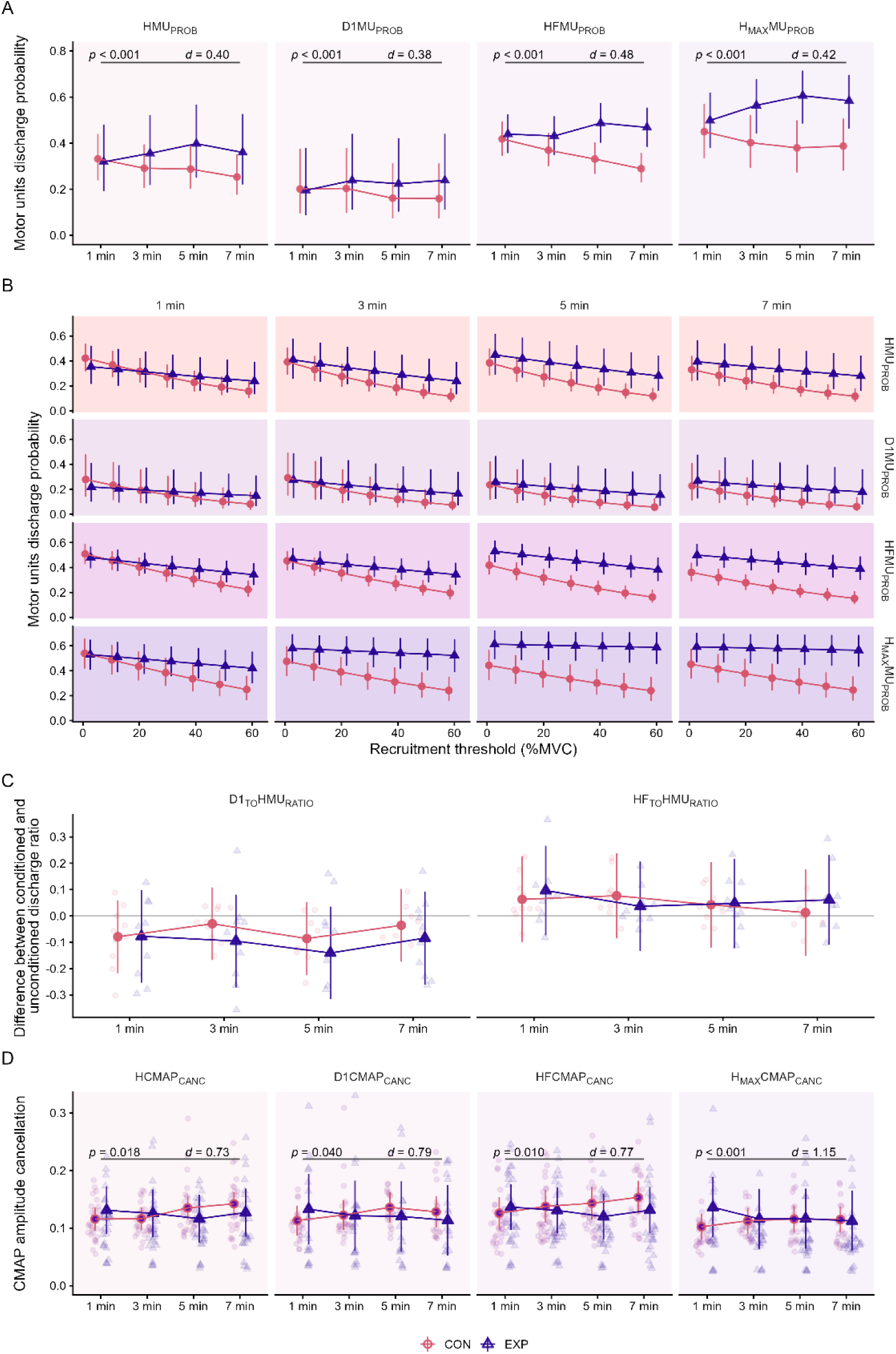
Motor unit (MU) discharges in elicited contractions. A) MU discharge probability. B) Graphical representation of the influence of recruitment threshold on MU discharge probability. C) Difference between conditioned and unconditioned discharge rate. D) CMAP amplitude cancellation. Statistical data are presented as estimated marginal means with 95% confidence intervals for both the control (CON, red circles) and experimental (EXP, blue triangles) conditions. Smaller and lighter colour circles and triangles represent individual raw data points in the background. Horizontal lines represent statistically significant interaction effects, accompanied by p values and Cohen d values above the relevant lines.

**Figure 7:**
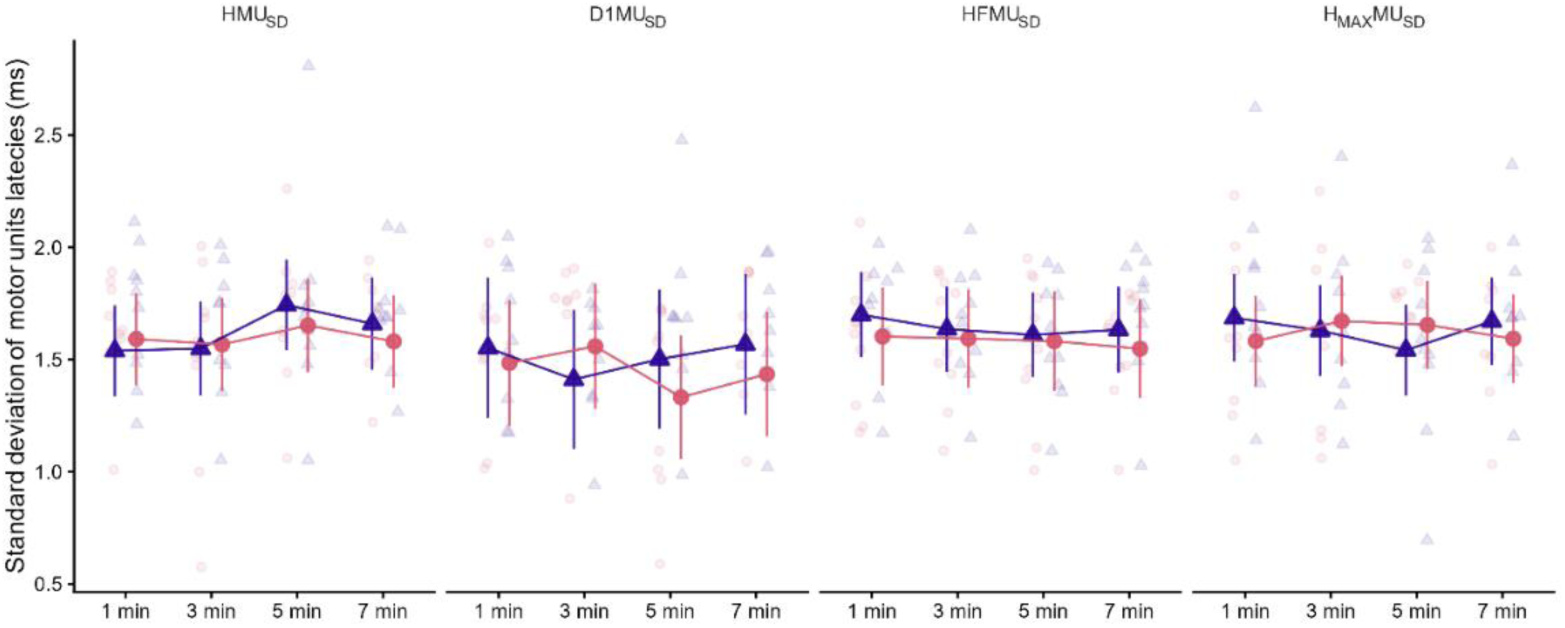
Mean standard deviation of MU firings in elicited contractions. Statistical data are presented as estimated marginal means with 95% confidence intervals for both the control (CON, red circles) and experimental (EXP, blue triangles) conditions. Smaller and lighter colour circles and triangles represent individual raw data points in the background.

MU discharge ratio was not significantly affected by the interaction between *time × intervention* in any of the assessed conditioning protocols (D1_TO_HMU_RATIO_: F_3 49.93_ = 1.18, p = 0.324; HF_TO_HMU_RATIO_: F _3 50.58_ = 1.26, p = 0.295).

CMAP amplitude cancellation for HCMAP_CANC_ and H_MAX_CMAP_CANC_ were significantly affected by the interaction between *time × intervention* (F_3, 205.33_ = 3.32, p = 0.020 and F_3, 157.19_ = 4.13, p = 0.007, respectively; Figure 6C). However, interaction contrasts showed no difference between interventions in any timepoint for HCMAP_CANC_ (Figure 6D), whereas H_MAX-CANC_ demonstrated a significant difference between 1 and 7 minutes (p = 0.008, d = 0.89). No significant *time × intervention* interaction was found for D1_CANC_ and HF_CANC_ (F_3, 163.68_ = 0.74, p = 0.524 and F_3, 224.18_ = 2.52, p = 0.058, respectively).

The *standard deviation* of MU discharges were not significantly affected by the interaction between *time × intervention* in any of the assessed conditioning protocols (HMU_SD_: F_3, 223.15_ = 0.49, p = 0.687; D1MU_SD_: F_3, 217.1_ = 2.17, p = 0.113; HFMU_SD_: F_3, 227_ = 0.25, p = 0.854; H_max_MU_SD_: F_1, 199.88_ = 1.42, p = 0.235).

## Discussion

This study aimed to investigate the immediate and short-term effects of PAP on neuromuscular responses following a maximal voluntary isometric conditioning contraction. Specifically, we assessed the muscle’s mechanical (single twitch) and neurophysiological (H-reflex) response, with the aim of elucidating the role of presynaptic spinal mechanisms. The main findings indicate that a brief maximal isometric conditioning contraction led to an increase in twitch torque, accompanied by a diminished H-reflex amplitude, which occurred despite indicators of reduced presynaptic inhibitory spinal mechanisms, as evidenced by increased D1 and HF responses. Modulation of mechanical and neurophysiological responses was transient in nature, vanishing after nine and three minutes after contraction, respectively. To the best of our knowledge, this is the first study to specifically examine spinal presynaptic mechanisms in the context of PAP.

### The pattern of mechanical and neurophysiological responses following a strong contraction

TW_PT_ was increased immediately after the conditioning contraction and persisted for about 9 minutes, demonstrating a strong potentiation effect, in line with previous studies investigating PAP (Baudry and Duchateau 2004; Folland et al. 2008; Wallace et al. 2019). Concomitantly, M_MAX_ remained unaffected by any intervention, consistent with a prior study (Folland et al. 2008), suggesting that sarcolemmal membrane excitability was not affected by the conditioning contraction (Rodriguez-Falces and Place 2021). Conversely, H_PTP_ was significantly decreased immediately following the conditioning contraction, but returned to baseline within three minutes. The decreased H_PTP_ immediately post-conditioning contraction is consistent with some studies (Enoka et al. 1980; Guellich and Schmidtbleicher 1996; Trimble and Harp 1998) but not others (Folland et al., 2008; Iglesias-Soler et al., 2011; Wallace et al., 2019), with the discrepancies between studies possibly attributed to variations in training status, conditioning contraction duration, intensity, type, and involved body region, as well as differences in the H-reflex methodology. Regarding the latter, studies interchangeably used the maximal H-reflex and H-reflex measured on the ascending part of the HM recruitment curve (Trimble & Harp (1998); like H_PTP_ used in this study). Indeed, in our study H_PTP_ significantly decreased, whereas the H_MAX_ remained statistically unchanged. While the lack of observed changes in H_MAX_ might also be due to the timing of the stimulation (i.e., the effect might have dissipated by the time H_MAX_ was assessed in our protocol), recent findings suggest that the maximal H-reflex may lack sensitivity to inhibitory mechanisms, potentially obscuring key physiological insights (Theodosiadou et al. 2022).

Prior studies have also reported an increase in reflex amplitude between 4 and 11 minutes after the contraction (Folland et al. 2008; Güllich and Sehmidtbleicher 2006), an effect not observed in our study. According to a recent review (Zero and Rice 2021) the delayed reflex potentiation reported in some studies might not be attributable to PAP, but to PAPE, a neuromechanical potentiation associated with mechanisms unrelated to PAP, such as increased muscle temperature, water content, and blood flow (Blazevich and Babault 2019). Furthermore, we can exclude changes in the onset of the reflex responses in the motoneuron or changes in the synchronisation of MU firings as the likely mechanisms for the observed H-reflex depression, due to no significant changes in unconditioned and conditioned H-reflex latencies (H_LAT_); (Knikou & Rymer, 2002) and no differences in the duration of the H-reflex (H_DUR_), respectively.

In previous studies, the drop in H_PTP_ immediately after conditioning contraction was suggested to be a compensatory mechanism to accommodate the enhanced muscular response (Zero and Rice 2021). However, the direct effect of enhanced muscle contractile properties to the spinal excitability is still unknown. Moreover, if the drop in H_PTP_ primarily reflects a compensatory mechanism, we would expect a return to baseline proportional to the drop in TW_PT_. In our data, the amplitude of H_PTP_ was inversely proportional to TW_PT_ only at the first timepoint. Thus, it is conceivable that other mechanisms contribute to the neural modulation of PAP.

### The role of presynaptic mechanisms in H-reflex reduction following a strong contraction

Contrary to our hypothesis, we observed significant increases in presynaptic inhibition-related parameters D1/H_PTP_ and HF/H_PTP_. Thus, changes of D1/H_PTP_ and HF/H_PTP_ suggest spinal facilitation or disinhibition rather than inhibition. This indicates a complex interplay between inhibitory and facilitatory spinal mechanisms following voluntary contractions and supports the idea that mechanisms other than presynaptic inhibition influence the observed reduction in H_PTP_. Observations from animal studies show that a short conditioning contraction elevates the transmittance of excitation potentials across synaptic junctions at the spinal cord (Gossard et al. 1994; Lüscher et al. 1983), which can explain the observed disinhibition/facilitation of D1/H_PTP_ and HF/H_PTP_. An induced tetanic contraction has been suggested to decrease the transmitter failure during subsequent activity, via one or a combination of several possible responses, including an increase in the quantity of neurotransmitter released, an increase in the efficacy of the neurotransmitter, or a reduction in axonal branch-point failure along the afferent neural fibres (Clamann et al. 1989; Lüscher et al. 1983). Moreover, consistent with our findings, following a conditioning contraction in the upper limbs that induced PAP, several studies reported reduced cervicomedullary motor evoked potential amplitude that has a large monosynaptic component and is generally not influenced by presynaptic inhibition (Jackson et al. 2006; Nielsen and Petersen 1994).

Another possible explanation for the drop in H_PTP_ observed in our study is the involvement of post-activation depression (PAD), a mechanism often attributed to the depletion of neurotransmitter release at the synaptic cleft due to previous activation of Ia afferents (Hultborn et al. 1996). PAD is known to be attenuated during sustained contractions, possibly due to enhanced Ia firing induced by the voluntary contraction (Burke 2016; Burke et al. 1989). In this context, the soleus and gastrocnemius lateralis H-reflex were found to be less depressed immediately after the conditioning contraction when assessed during a sustained voluntary contraction compared to rest, which was interpreted as evidence of PAD-induced depression of the H reflex amplitude (Xenofondos et al. (2015)).

The transient reduction in H_PTP_ observed in the present study could also be explained by a rapid, history-dependent decrease in the sensitivity of muscle spindle Ia afferents, a peripheral phenomenon known as muscle thixotropy (Proske and Morgan 1999). In our protocol, the strong conditioning contraction likely caused the soleus muscle and its spindles to shorten. Upon relaxation, the muscle fibres, having formed stable cross-bridges at a shortened length, would be in a slack state at the testing position. While the presence of slack in the whole muscle may not always be apparent, the muscle spindles with their compliant connections to adjacent extrafusal fibres are particularly prone to slack (Proske et al. 1993), which dramatically reduces tension on the spindle’s sensory endings, lowering their baseline discharge rate. This is supported by data in cats, where a conditioning contraction has been shown to dramatically reduce discharge rate of primary spindle endings (from 40 to 10 pps), suggesting a profound, momentary desensitization of the sensory part of the muscle spindle without any change in actual muscle length (Gregory et al. 1986). In the case of our study, the standardised submaximal stimulus that activates the axons originating from spindles will have evoked a less synchronous or less potent volley of action potentials arriving at the spinal cord, leading to weaker excitatory post-synaptic potential generated by the alpha motoneuron when the spindle axons will have been in a thixotropic, low-sensitivity state after a conditioning contraction, resulting in decreased H-reflex. Indeed, the human evidence supports this supposition, showing that conditioning contractions that alter muscle thixotropy can modulate H-reflex amplitude (Wood et al. 1996). Although postsynaptic mechanisms such as post-activation depression represent a possible explanation for the apparent contradiction in in our data (i.e. decreased H-reflex with presynaptic inhibition measures pointing toward facilitation), thixotropy offers an additional plausible explanation where the decreased H-reflex reflects a peripheral decrease in Ia-afferent efficacy rather than a change in spinal inhibition. Future studies designed to directly assess post activation depression and presynaptic inhibition while manipulating muscle thixotropy (e.g., via specific conditioning contractions) are needed to test this hypothesis explicitly.

### Insights on the effects of PAP on spinal mechanisms gleaned from single motor units

Additional insights into the effects of PAP on spinal mechanisms were obtained through the decomposition of the HDsEMG signals into contributions of individual MUs. MU discharge latency was not affected by the conditioning contraction, confirming the results observed from global EMG metrics. Moreover, the MU discharge probability was similar between conditions immediately after the conditioning contraction. However, discharge probability increased three minutes after the contraction in the EXP compared to the CON condition across all stimulation paradigms and was significantly higher in higher-threshold MUs. This observation is in line with animal studies where tetanic contraction decreased the transmitter failure occurring primarily at larger motoneurons, which resulted in a considerable potentiation effect at this motoneurons (Lüscher et al. 1983). In addition, CMAP amplitude cancellation, a measure of MU discharge synchronisation, was lower in the EXP condition, suggesting increased synchronisation of MU discharges following the conditioning contraction. Overall, the combination of lower CMAP amplitude cancellation and higher discharge probability, particularly in higher-threshold units, suggests that the conditioning contraction induced facilitation rather than inhibition. These results contradict the reflex amplitude metrics observed at the global EMG level (H_PTP_), not only in the direction of change but also in timing. Specifically, MU-level differences emerged three minutes after the contraction, whereas the decrease in H_PTP_ occurred immediately. Previous studies have suggested that PAP may increase MU recruitment while decreasing MU discharge rate (Inglis et al. 2011; Klein et al. 2001), which could partially explain this discrepancy.

When interpreting the discrepancy between single MU variables and global EMG metrics reported in this study, it is important to note that MU discharge probability metrics do not represent a fair comparison to D1_PTP_/H_PTP_ and HF_PTP_/H_PTP_ which are normalized to the H_PTP_ amplitude. Thus, we computed the D1_TO_H_RATIO_ and HF_TO_H_RATIO_, where we subtracted the number of discharged MUs in the unconditioned contraction from D1 and HF, respectively, and expressed this difference as a ratio with respect to the total identified MUs of each participant. The analysis of the discharge ratio shows that analysis at the MU level also captures the pattern of D1 inhibition and HF facilitation, which can be seen as a decrease or increase in D1_TO_H_RATIO_ and HF_TO_H_RATIO_, respectively. However, discharge ratio was not sufficiently sensitive to capture the increase in D1 and HF, as seen in the global EMG (D1_PTP_/H_PTP_ and HF_PTP_/H_PTP_). In this respect, it is important to acknowledge that HDsEMG decomposition captures only a subset of active MUs, with a bias toward larger and more superficial units (Farina et al. 2010), whereas global EMG provides a more integrated representation of the whole muscle (Vieira and Botter 2021). Furthermore, amplitude metrics from global EMG are continuous and non-affected by single-unit errors, whereas MU-level analyses rely on a relatively small number of units and discharges. As such, mislabelling or missing a few MU discharges can disproportionately influence the results. Though the single MU analysis provided complementary evidence to increased spinal excitability following voluntary contraction particularly in higher-threshold units, future studies employing a higher stimulus count are warranted to fully exploit the potential of this method for detailed mapping of motoneuron pool adaptations.

### Limitations

This study presents several limitations. First, although the stimulation block intended to induce the H-reflex in the ascending part of the H/M recruitment curve, the D1, HF, and maximal H reflex blocks lasted 90 seconds, and only four stimuli were elicited per response. While 5–10 reflexes are typically recommended for reliable assessment (Theodosiadou et al. 2022), the approach used in this study allowed us to examine multiple neurophysiological mechanisms within a single session with the low variability within each time point (Figure 4A) suggesting four stimuli were sufficient to capture the modulation.

Second, although similar intervals were used in prior work (Iglesias-Soler et al. 2011), we used a shorter interstimulus interval (6 seconds) than the 10 seconds typically recommended for resting H-reflex assessments to minimize post-activation depression (Palmieri et al. 2004). Varying intervals were tested during the pilot and familiarization phases, allowing us to identify the shortest interval that did not induce post-activation depression, thereby increasing the number of stimuli per block. Maximal twitches were elicited immediately after the conditioning contraction and could theoretically influence subsequent H reflexes. However, an identical stimulation paradigm was used in the CON condition, which did not show any decrease in H reflex amplitude, suggesting that prior maximal stimulations did not bias the reflex responses. The maximal H reflex was assessed 60 seconds after H_PTP_, which may have allowed PAP-induced modulation to dissipate. Nonetheless, HF assessed just 20 seconds earlier remained significantly elevated, suggesting that the absence of H_MAX_ modulation more likely reflects a methodological limitation of H_MAX_, rather than an absence of neurophysiological effects.

Lastly, MU-level analysis is biased by the small number of units identifiable within each evoked response. Although our group has previously demonstrated the feasibility of extracting MUs from evoked responses (Kalc et al. 2022a, 2022b; Škarabot et al. 2023), achieving physiologically robust results still requires hundreds of responses. For example, in a recent study (Magdič et al. 2025) we successfully used PSTHs to extract information from the earliest portion of the reflex response, which reflects the monosynaptic phase of Ia presynaptic inhibition (Hultborn et al. 1987). Although this analysis reinforced the physiological effects observed at the global EMG level, a greater number of evoked responses than the number used in the present study is required for HDsEMG-derived MU metrics to reach the reliability comparable to intramuscular EMG. Future attempts to apply similar techniques to study fast, transient neurophysiological phenomena such as PAP should consider incorporating low-intensity background muscle activity to reduce post-activation depression (a common limitation in resting H-reflex paradigms) and enable the acquisition of a much greater number of stimuli within a similar assessment window. This approach would strengthen the robustness of non-invasive spinal assessments and enhance the mechanistic insights available from PSTH/PSF analyses.

### Conclusions

This study offers novel insights into the neuromuscular responses to post-activation potentiation following a maximal voluntary isometric conditioning contraction, particularly regarding spinal mechanisms. Our results indicate that PAP, induced by a brief but intense voluntary contraction, enhances twitch torque, accompanied by a disinhibition (facilitation) of presynaptic inhibitory mechanisms, as shown by increased D1 and HF responses. The observed reduction in H reflex amplitude immediately after the contraction, however, suggests that additional spinal mechanisms or an intrinsic muscle contraction history-dependent mechanical state may modulate spinal output. Future studies should therefore dissociate spinal mechanisms from muscle thixotropy using protocols that isolate their independent effects.

## Supporting information

S1 contains the inpact of numbe of discharging MUs on CMAP and our mitigation

