## Supplementary material for "Spinal Mechanisms in Post-Activation Potentiation: Facilitation of Presynaptic Inhibition Contrasts H-Reflex Amplitude Reduction": S1 contains the inpact of numbe of discharging MUs on CMAP and our mitigation

**Supplementary Material: Rationale and Methodology for Exclusion of Data Points in Cancellation Analysis**

In this supplementary document, we detail the rationale and methods used in the processing of EMG cancellation, with particular emphasis on the criteria used for excluding specific data points from the final analysis.

Figure S1 illustrates the CMAP cancellation values in relation to the number of MU discharges. For each stimulation condition (unconditioned H-reflex, D1 presynaptic inhibition, HF homonymous facilitation, and maximal H-reflex). Each datapoint represents a single reflex response. A cancellation value of 1 indicates the lowest possible cancellation (highest peak-to-peak amplitude of the CMAP) and in our study occurs when only a single MU has been detected, a case in which cancellation analysis is physiologically meaningless because cancellation quantifies temporal dispersion across multiple firings. As visible in the Figure S1, several responses cluster at this upper bound, reflecting insufficient MU discharges for cancellation analysis. Moreover, responses with few detected MU discharges (below 10) tend to produce disproportionately high cancellation values, representing a methodological artefact due to an insufficient amount of data rather than genuine physiological synchrony. Thus responses with 10 and more firing MUs were used in the statistical analysis of CMAP cancellation.

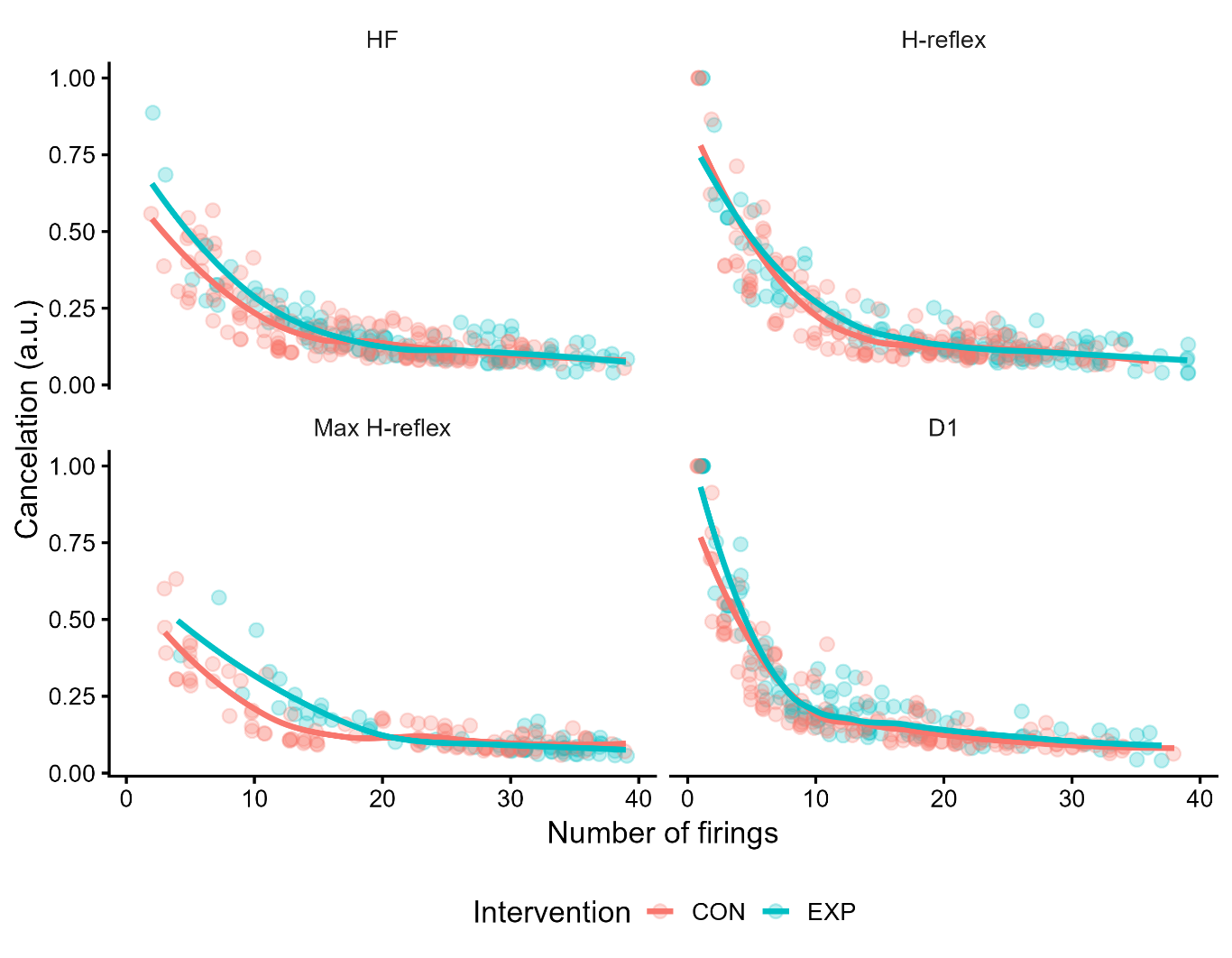

Figure S1: Raw CMAP cancellation values for the first four timepoints following the intervention (1 to 7 minutes) across all stimulation conditions: unconditioned H-reflex, D1-conditioned H-reflex, HF-conditioned H-reflex, and maximal H-reflex. Each datapoint represents a single response.
